# A robust and self-sustained peripheral circadian oscillator reveals differences in temperature compensation properties with central brain clocks

**DOI:** 10.1101/2020.06.24.168450

**Authors:** Marijke Versteven, Karla-Marlen Ernst, Ralf Stanewsky

## Abstract

Circadian clocks temporally organize physiology and behavior of organisms exposed to the daily changes of light and temperature on our planet, thereby contributing to fitness and health. Circadian clocks and the biological rhythms they control are characterized by three properties. (1) The rhythms are self-sustained in constant conditions with a period of ~ 24 hr, (2), they can be synchronized to the environmental cycles of light and temperature, and (3), they are temperature compensated, meaning they run with the same speed at different temperatures within the physiological range of the organism. Apart from the central clocks located in or near the brain, which regulate the daily activity rhythms of animals, the so-called peripheral clocks are dispersed throughout the body of insects and vertebrates. Based on the three defining properties, it has been difficult to determine if these peripheral clocks are true circadian clocks. We used a set of clock gene – luciferase reporter genes to address this question in *Drosophila* circadian clocks. We show that self-sustained fly peripheral oscillators over compensate temperature changes, i.e., they slow down with increasing temperature. This over-compensation is not observed in central clock neurons in the fly brain, both in intact flies and in cultured brains, suggesting that neural network properties contribute to temperature compensation. However, an important neuropeptide for synchronizing the circadian neuronal network, the Pigment Dispersing Factor (PDF), is not required for self-sustained and temperature-compensated oscillations in subsets of the central clock neurons. Our findings reveal a fundamental difference between central and peripheral clocks, which likely also applies for vertebrate clocks.

## Introduction

Circadian clocks temporally organize physiology and behavior of all organisms exposed to the daily changes of light intensities and temperature. There are growing numbers of examples that circadian clock function positively contributes to an organism’s fitness and to human health, highlighting the importance of understanding their underlying molecular mechanisms. In order to function as reliable time keepers, one hallmark of circadian clocks is their temperature independence (e.g., (Hall, 1997; Pittendrigh, 1954)). Particularly in poikilothermic animals, a circadian clock that would change its pace with temperature would be of little use to accurately measure time. As a consequence, circadian clocks are ‘temperature compensated’ meaning they keep ticking with a period close to 24 hr over a large variation of ambient temperatures within the organism’s physiological range (Hall, 1997). This ability to keep a temperature-compensated 24-hr rhythm therefore is also one of the three characteristics that define a circadian rhythm, the other two being the self-sustained nature and the ability to be entrained by environmental and other time cues (Zeitgeber).

In *Drosophila*, the free-running period length (Tau, τ) for both rhythmic eclosion of pharate adults from the pupal case, as well as adult locomotor activity, remains almost exactly 24 hr between 16°C and 28°C, revealing a remarkable degree and precision of temperature compensation (Konopka et al., 1989; Pittendrigh, 1954; Singh et al., 2019; Zimmerman et al., 1968). Both eclosion and locomotor activity rhythms are controlled by specialized clock neurons in the pupal or adult brain (Ewer et al., 1992; Myers et al., 2003). These clock neurons are defined through their expression of clock genes like *period* (*per*), *timeless* (*tim*), *Clock* (*Clk*) and *cycle* (*cyc*), which are all required to maintain the rhythmic behaviors of the fly (Peschel and Helfrich-Förster, 2011). These clock genes form the molecular bases of the circadian clock by generating self-sustained 24-hr oscillations of clock gene transcription and protein abundance. Transcription is activated by binding of the transcription factors CLK and CYC to *per* and *tim* promoter elements. PER and TIM proteins slowly accumulate in the cytoplasm before entering the nucleus, where they shut down their own transcription by binding to CLK and CYC. Only after PER and TIM degradation this 24-hr lasting cycle can start again. The period length of this molecular cycle is controlled by the stability and subcellular localization of the PER and TIM repressor proteins, which are both determined by several kinases and phosphatases targeting PER and/or TIM (Tataroglu and Emery, 2015). Considering the molecular and biochemical nature of this feedback loop underlying circadian clocks, their ability to drive temperature compensated biological rhythms seems surprising. Current models predict that counter balancing biochemical reactions, some that speed up, other that slow down with increasing temperature, result in overall temperature independent circadian oscillations (e.g., (Shinohara et al., 2017)).

Based on the temperature independent τ of clock controlled behaviors, it seems required that the molecular oscillations and activity of the central clock neurons driving these output rhythms must somehow be temperature compensated. This is less clear for peripheral clocks for several reasons. Peripheral clocks are widely distributed within the fly, for example in eyes, wings, legs, antennae, proboscis, Malpighian tubules, testes, and even the cuticle (Beaver et al., 2002; Chatterjee et al., 2010; Ito et al., 2008; Ivanchenko et al., 2001; Levine et al., 2002; Plautz et al., 1997). Yet, only for the antennal, proboscis, and cuticle peripheral clocks it is known which biological output rhythm they control, but it is not known if these rhythms are temperature compensated (Chatterjee et al., 2010; Ito et al., 2008; Krishnan et al., 1999). In addition, the molecular machinery in peripheral clocks differs from that in central clock neurons. The main difference concerns the function of *cryptochrome* (*cry*), which encodes a blue-light photoreceptor for the synchronization of both central clock neurons and peripheral clocks (Ito et al., 2008; Ivanchenko et al., 2001; Stanewsky et al., 1998). In addition, CRY appears to be required for clock function in most peripheral tissues, for example in the eye and the antennae (Krishnan et al., 2001; Levine et al., 2002; Stanewsky et al., 1998) presumably by acting as repressor for *per* and *tim* expression, similar to the mammalian CRY proteins (Collins et al., 2006). Although peripheral clocks can be entrained to light:dark (LD) and temperature cycles (Glaser and Stanewsky, 2005; Harper et al., 2017; Ito et al., 2011; Ivanchenko et al., 2001; Levine et al., 2002), it surprisingly is still unclear if they meet the remaining two criteria, which would qualify them as genuine circadian clocks: self-sustained rhythmicity and temperature compensation. It is difficult to determine if peripheral clocks are temperature compensated, because their molecular oscillations dampen out very quickly in constant conditions, mitigating against reliable estimation of τ (e.g. (Harper et al., 2017; Kidd et al., 2015; Levine et al., 2002; Veleri et al., 2003)). Furthermore, it is not clear if this dampening is due to the lack of a self-sustained molecular oscillator or caused by rapid desynchronization between individual peripheral oscillator cells. To address the properties of peripheral clocks in more detail, we applied an established luciferase reporter system, allowing to measure clock gene expression in real time in live tissues (e.g. (Glaser and Stanewsky, 2005)). We identified a robust, self-sustained 24-hr oscillator in the fly’s haltere, an organ important for proprioceptive feedback during flight and in some insects also walking (Yarger and Fox, 2016). Surprisingly, the haltere clock is over-compensated against temperature changes, i.e., it slows down with increasing temperature rather than speeding up. We show that the same applies for the antennal peripheral clock, indicating that this a general feature of peripheral fly oscillators. Using the same assay, we show that molecular oscillations in the central brain clock are perfectly temperature compensated, neither slowing down nor speeding up with temperature. While this difference indicates the importance of neuronal network properties, we show that the neuronal synchronizing peptide PDF is not required for accurate temperature compensation.

## Results

### The *Drosophila* haltere contains a robust and self-sustained peripheral circadian oscillator

In contrast to circadian pacemaker neurons in the fly brain, peripheral circadian clocks of *Drosophila* are generally considered as weak or dampened circadian oscillators, due to their inability of maintaining molecular oscillations in constant conditions (e.g., (Levine et al., 2002; Stanewsky et al., 1997; Veleri et al., 2003). We were therefore surprised to observe sustained and non-dampening oscillations during constant conditions in cultured halteres of transgenic *ptim-TIM-luc* flies, a reporter for TIM protein expression ((Lamba et al., 2018), Figures 1A, B, S1). After initial synchronization to LD cycles, bioluminescence signals from individual pairs of halteres were measured in constant darkness and temperature (DD, 25°C) with 30 min or 1 hr time resolution. In order to determine the period-length of the bioluminescence oscillations, data were detrended and subjected to curve fitting (Figure 1C, (Klemz et al., 2017)). At 25°C, *ptim-TIM-luc* oscillations have a τ of 24.0 ± 0.1 hr demonstrating that they are indeed circadian (Figure 1D, Table 1). In addition, we analyzed *ptim-TIM-luc* expression in halteres of clock-less *per*^*01*^ flies, which lead to arrhythmic reporter gene expression as expected (Figure S2).

**Table 1:** Free running period (τ) and expression levels of clock gene expression in halteres, and clock neurons. τ values were calculated in Chronostar (see Transparent Methods). The ‘Error’ value, depicts the correlation between the curve fit and the detrended data (1 minus correlation coefficient, so the lower the error value the more trustable the period value). All individual τ values with an error <0.5 or <0.7 were used for calculating the average τ in the tissue culture, and adult fly experiments, respectively. Q_10_ values were calculated using the τ values for 18°C and 29°C. The ‘mean’ reflects the average expression level (CPS) during the entire time series. The brain *tim-TIM-luc* LV200 values refer to the two DN1 groups recorded at 21°C with the LV200 CCD camera system (Figure 4C-F and Movie S1).

| Genotype | °C | N | Tau ± S.E.M (h) | Error ± S.E.M | Mean ± S.E.M (CPS) |
| --- | --- | --- | --- | --- | --- |
| **halteres** |  |  |  |  |  |
| *plo* | 18 | 23 | 22.8 ± 0.2 | 0.15 ± 0.01 | 377.9 ± 31.3 |
| | 21 | 30 | 22.9 ± 0.1 | 0.04 ± 0.01 | 302.4 ± 18.9 |
| | 25 | 28 | 24.2 ± 0.1 | 0.04 ± 0.01 | 385.2 ± 35.7 |
| | 29 | 11 | 24.4 ± 0.2 | 0.05 ± 0.01 | 190.2 ± 23.1 |
|  |  |  | **$Q_{10} = 0.94$** |  |  |
| *tim-luc* | 18 | 25 | 22.1 ± 0.2 | 0.32 ± 0.03 | 505.6 ± 33.7 |
| | 21 | 23 | 23.0 ± 0.1 | 0.07 ± 0.01 | 705.7 ± 53.4 |
| | 25 | 33 | 24.6 ± 0.1 | 0.03 ± 0.01 | 1361.8 ± 85.8 |
| | 29 | 25 | 24.8 ± 0.1 | 0.03 ± 0.01 | 1340.3 ± 85.5 |
|  |  |  | **$Q_{10} = 0.90$** |  |  |
| *XLG-luc* | 18 | 22 | 19.9 ± 0.4 | 0.25 ± 0.03 | 115.3 ± 6.1 |
| | 21 | 12 | 20.2 ± 0.3 | 0.26 ± 0.03 | 117.4 ± 9.7 |
| | 25 | 28 | 21.2 ± 0.1 | 0.07 ± 0.01 | 157.2 ± 7.2 |
| | 29 | 14 | 23.4 ± 0.1 | 0.07 ± 0.01 | 149.1 ± 8.6 |
|  |  |  | **$Q_{10} = 0.86$** |  |  |
| *ptim-TIM-luc* | 18 | 27 | 21.0 ± 0.1 | 0.17 ± 0.02 | 77.7 ± 4.2 |
| | 21 | 37 | 21.7 ± 0.1 | 0.09 ± 0.01 | 99.2 ± 5.8 |
| | 25 | 53 | 24.0 ± 0.1 | 0.10 ± 0.01 | 112.2 ± 8.6 |
|  | 29 | 67 | 25.2 ± 0.1 | 0.04 ± 0.01 | 134.4 ± 6.9 |
|  |  |  | **$Q_{10} = 0.85$** |  |  |
| **brains** |  |  |  |  |  |
| *ptim-TIM-luc* | 18 | 14 | 28.6 ± 0.4 | 0.33 ± 0.02 | 184.3 ± 16.6 |
| TopCount | 21 | 13 | 30.8 ± 0.6 | 0.20 ± 0.02 | 167.7 ± 18.2 |
|  | 25 | 14 | 30.0 ± 0.8 | 0.30 ± 0.03 | 187.7 ± 8.1 |
|  | 29 | 5 | 30.0 ± 1.2 | 0.32 ± 0.06 | 324.3 ± 80.1 |
|  |  |  | **$Q_{10} = 0.96$** |  |  |
| LV200 | 21 | 1 | 31.7 | 0.18 | N/A |
|  | 21 | 1 | 31.4 | 0.20 | N/A |
| **flies** |  |  |  |  |  |
| *8.0-luc* | 18 | 38 | 24.8 ± 0.3 | 0.42 ± 0.03 | 85.1 ± 5.2 |
| *Cut off < 0.7* | 21 | 10 | 23.9 ± 0.8 | 0.59 ± 0.04 | 89.9 ± 10.4 |
|  | 25 | 31 | 24.5 ± 0.2 | 0.43 ± 0.03 | 71.5 ± 5.8 |
|  | 29 | 17 | 23.2 ± 0.3 | 0.17 ± 0.02 | 40.3 ± 2.9 |
|  |  |  | **$Q_{10} = 1.06$** |  |  |
| *8.0-luc; Pdf<sup>01</sup>* | 18 | 41 | 21.7 ± 0.2 | 0.27 ± 0.03 | 23.1 ± 0.9 |
| *Cut off < 0.7* | 21 | 18 | 22.7 ± 0.1 | 0.54 ± 0.03 | 17.7 ± 0.7 |
|  | 25 | 30 | 21.7 ± 0.1 | 0.27 ± 0.03 | 20.4 ± 1.0 |
|  | 29 | 15 | 22.5 ± 0.1 | 0.26 ± 0.03 | 15.2 ± 0.8 |
|  |  |  | **$Q_{10} = 0.97$** |  |  |

**Figure 1:**
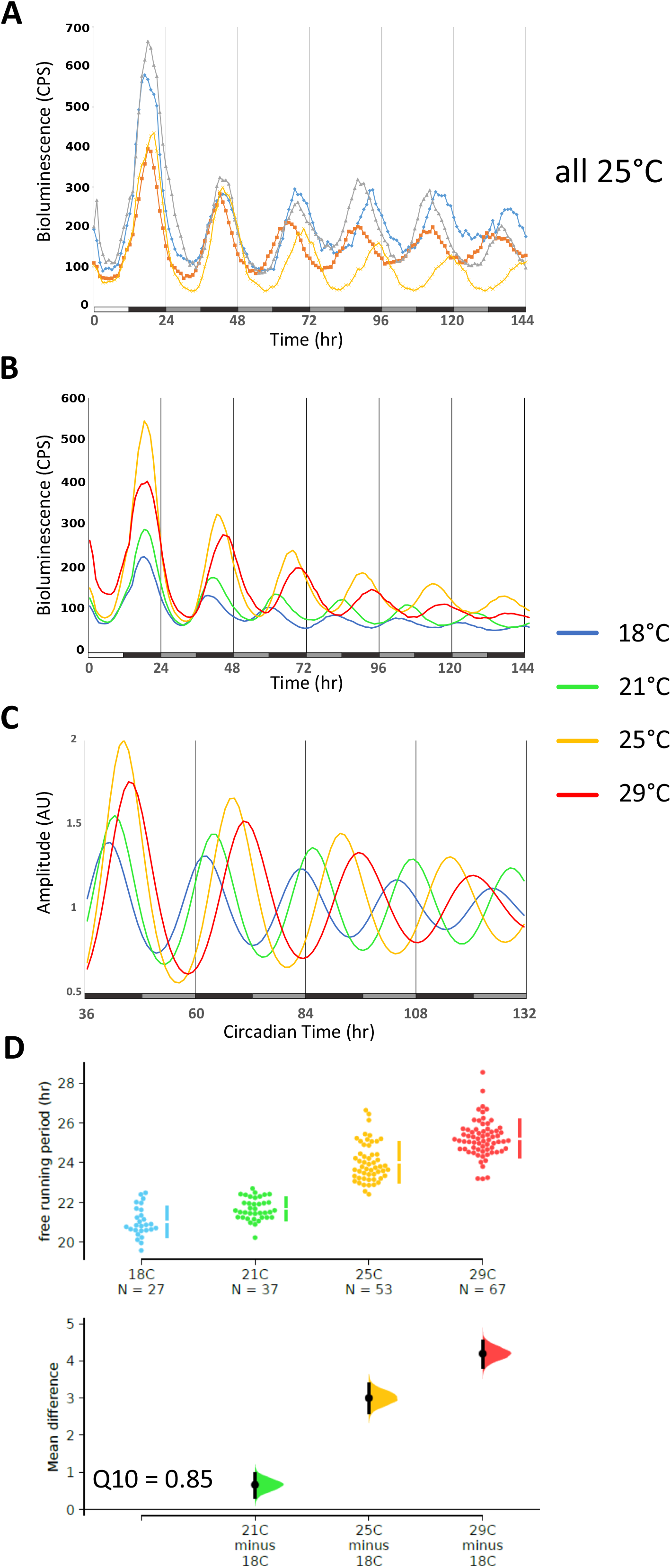
The *Drosophila* haltere contains a non-dampened circadian oscillator. Bioluminescence recordings of halteres dissected from *tim-TIM-luc* flies. Tissues were exposed to 2 LD cycles (data of the 1^st^ LD cycle were discarded) followed by DD at different constant temperatures. **(A)** Raw data traces of 4 pairs of halteres at 25°C. White portion of x-axis bar indicates ‘lights on’, black portion ‘lights-off’, and grey portions subjective day (= lights-off, but the time lights would have been on if the cultures were still in LD). CPS: counts per second. **(B)** Average traces at 18°C (n=27), 21°C (n=12), 25°C (n=53), 29°C (n=67), same conditions as in (A). **(C)** Curve-fitted oscillations after detrending the DD data in (B). **(D)** Period estimates for individual haltere pairs obtained from curve-fitted data (see Transparent Methods). To precisely visualize potential period differences between temperatures we applied estimation statistics (ES) rather than significance tests (Ho et al., 2019). Because ES focusses on the magnitude of the effect size (i.e., mean difference) and its precision, rather than acceptance or rejection of the null hypothesis, ES gives a more informative way to analyze and interpret results (Ho et al., 2019). We generated shared-control plots, which are analogous to an ANOVA with multiple comparisons, whereby the 18°C data served as shared-control. The top part shows all the period and expression level data points observed, with the mean and standard error plotted as a discontinuous line to the right (the gap indicates the observed mean). The bottom part a difference axis displays the effect size (mean difference, the black dots). The colored filled curves indicate the resampled distribution, and the black vertical line the 95% confidence interval (CI). The size of the CI shows the precision of the mean difference: if it crosses the reference line (x-axis) both datasets may originate from the same distribution (p > 0.05), if not, they are different (p < 0.05) (see Transparent Methods and (Ho et al., 2019) for more details). See also Table 1 for numeric τ and expression level values.

### Clock protein oscillations in the haltere slow down with increasing temperature

We next asked, if these oscillations are also temperature compensated, as this feature is a defining criterion for circadian clocks (Pittendrigh, 1954). To our surprise, although robust oscillations were also observed at 18°C, 21°C, and 29°C, τ increased quite dramatically by 4.2 hr from the lowest (18°C) to the highest (29°C) temperature (Q_10_ = 0.85) (Figure 1B-D, Table 1). These results indicate that the haltere circadian oscillator is temperature compensated, because it does not speed up with increasing temperature. Nevertheless, this temperature compensation is not normal, since the oscillations slow down with temperature, therefore suggesting an ‘over-compensation’ against temperature changes. In order to investigate if this over- compensation is specific for the TIM protein, we next performed the same experiments with *XLG-luc* transgenic flies, expressing a reporter gene that reflects spatial and temporal PER expression (Veleri et al., 2003) (Figure S1). Comparable to the *ptim-TIM-luc* reporter, *XLG-luc* halteres showed robust oscillations at all 4 temperatures (Figures 2A, S3A). Similar to what we observed for *ptim-TIM-luc*, the τ associated with the *XLG-luc* oscillations increased by about 3.5 hr with increasing temperature (Q_10_ = 0.86), suggesting that the oscillations of clock proteins in halteres are over-compensated against temperature changes (Figures 2A, S3A, Table 1).

**Figure 2:**
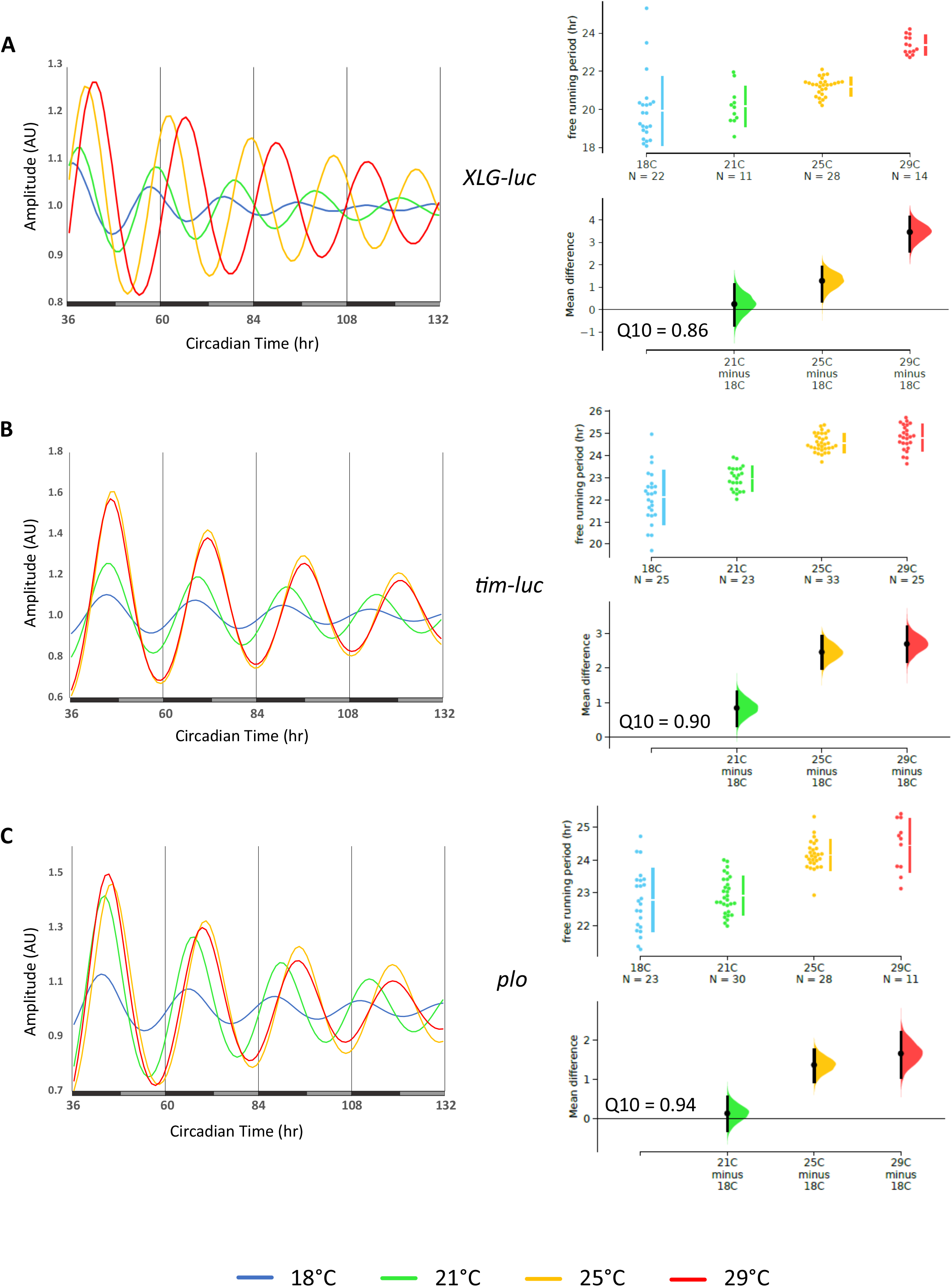
Clock protein rhythms in the haltere slow down with increasing temperature. Bioluminescence recordings of halteres dissected from **(A)** *XLG-luc*, **(B)** *tim-luc*, and **(C)** *plo* at the indicated constant temperatures in DD. The plots on the left show curve-fitted data for the DD portion of the experiment. To the right, ES shared control plots comparing the distribution of period values at the different temperatures (see legend to Figure 1D and Transparent Methods for details).

To investigate if the temperature compensation phenotype in halteres extends to clock gene transcription, we employed transcriptional reporters for *per* (*plo*) (Brandes et al., 1996) and *tim* (*tim-luc*) (Stanewsky et al., 2002) (Figure S1) and performed the same experiments as described above for the protein reporters. Halteres expressing *tim-luc* were robustly rhythmic at all 4 temperatures (Figures 2B, S3A). Interestingly, τ increased only by 2.7 hr from 18°C to 29°C (Q_10_ = 0.90), indicating less over-compensation of *tim* transcription compared to PER and TIM proteins (Figures 2B, S3A, Table 1). Similar results were obtained for *plo* halteres, which exhibited high amplitude oscillations at all 4 temperatures with a temperature-dependent τ increase of only 1.6 hr (Q_10_ = 0.94) (Figures 2C, S3, Table 1). In summary, the results show that PER and TIM oscillations in halteres are over-compensated, largely caused by posttranscriptional mechanisms leading to period-lengthening with increasing temperatures.

In addition to temperature-dependent changes in τ, we also observed differences in the expression levels of the same transgene at different temperatures. Consistent with previous observations (Kidd et al., 2015; Majercak et al., 1999), both PER and TIM protein reporters show higher expression at 25°C and 29°C compared to the lower temperatures (Figure S3, Table 1). From these results it could be expected that due to the repressor activity of PER and TIM, transcription rates of *per* and *tim* would be lower at the warm temperatures compared to the cooler ones. Indeed, in the case of *per*, low protein (*XLG-luc*) expression levels at 18°C and 21°C compared to 25°C and 29°C, coincide with elevated levels of *per* transcription (*plo*) at 18°C and 21°C compared to 29°C (Figure S3, Table 1). In contrast, expression levels of both *tim* constructs were significantly reduced at 18°C compared to the higher temperatures, whereas there was no difference (*tim-luc*) or a slight increase (*tim-TIM-luc*) between 25°C and 29°C (Figure S3; Table 1). These results are not readily compatibly with direct negative feedback regulation through PER and TIM repression, where one would expect higher protein levels to be correlated with a lower rate of *tim* transcription.

### Peripheral clocks in the antennae slow down with increasing temperatures

To investigate if overcompensation against temperature changes is a general feature of *Drosophila* peripheral clocks, we next performed the same analysis of clock gene expression in dissected antennae. This sensory organ contains a circadian clock controlling the daily sensitivity changes of the olfactory system (Krishnan et al., 1999). Consistent with published reports (Krishnan et al., 2001; Levine et al., 2002), rhythmic gene expression was readily detectable in DD at 25°C after initial synchronization to LD cycles in antennal pairs prepared from the four different luciferase transgenic strains (Figure 3, S4A, Table S1). In contrast to halteres, robust rhythmic reporter gene expression was not observed at all temperatures (in particular at 29°C), indicating that the antennal oscillator is less robust at least in warmer temperatures (Figures 3, S4A, Table S1). We therefore decided to determine temperature compensation properties of the antennal oscillator only for 18°C, 21°C, and 25°C, where we obtained reliable period estimates (Figures 3, S4A, Table 1). Comparing *ptim-TIM-luc* period values at the 3 different temperatures revealed a severe over-compensation phenotype, with a lengthening of τ by > 4 hr (Q_10_ = 0.79) (Figures 3, S4A, Table S1). In contrast, *XLG-luc* antennae, which were arrhythmic at 29°C, lengthened their period only by > 1 hr between 18°C and 25°C (Q_10_ = 0.92) (Figure 3, S4A, Table S1), indicating less over-compensation for this reporter. Nevertheless, the fact that τ increases significantly at 25°C compared to 18°C and 21°C and because expression becomes arrhythmic at 29°C, may indicate a strong over-compensation phenotype of PER expression in antennae, resulting in arrhythmicity instead of severe period lengthening. Strikingly, and in contrast to the situation in halteres, gene expression rhythms of the two transcriptional reporters for *per* and *tim* also showed severe over-compensation with a dramatic τ lengthening > 5-6 hours between 18°C and 25°C (Q_10_ = 0.77 and 0.69, respectively) (Figures3; S4A, Table S1).

**Figure 3:**
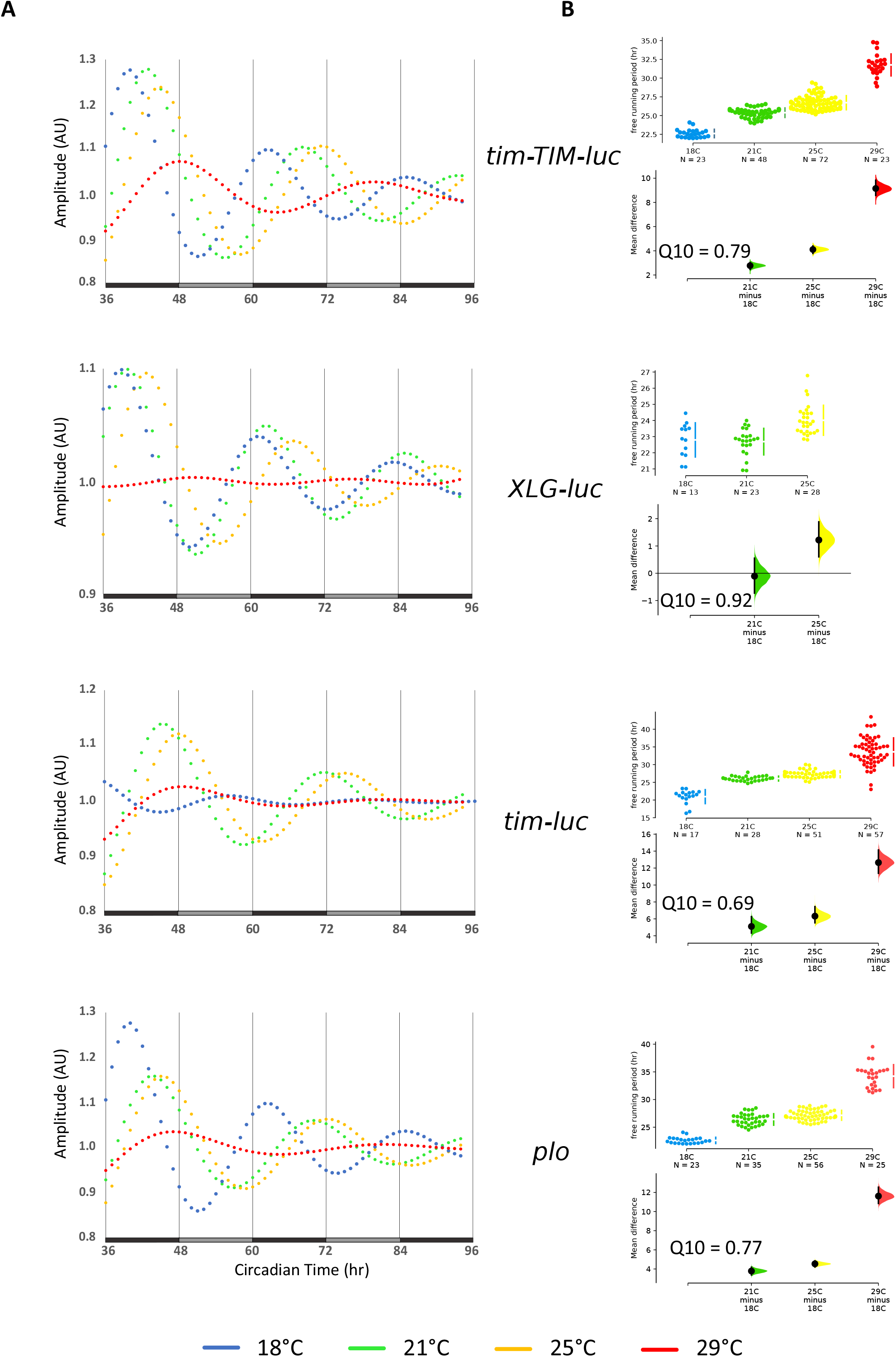
The antennal clock slows down with increasing temperature. **(A)** Curve fits of bioluminescence recordings of antennae dissected from *tim-TIM-luc*, *XLG-luc*, *tim-luc*, and *plo* flies at the indicated constant temperatures in DD. Black bars indicate subjective night, grey bars subjective day. **(B)** Period estimates for antennal pairs obtained from curve-fitted data in (A).

Mean expression levels of all four reporters changed with temperature in a very similar manner, with low expression levels at the extremes (18°C and 29°C), and higher expression levels at 21°C and 25°C (Figure S4, Table S1), precluding any meaningful correlation between protein levels and transcription rates as was the case for the halteres. It has previously been reported that luciferase activity is temperature-dependent, either increasing or decreasing with temperature (e.g. (Kurosawa et al., 2017; McElroy and Seliger, 1961)). Although we cannot rule out direct effects of temperature on luciferase activity, we do not observe consistent changes of luciferase activity with temperature. For example, *tim-luc* bioluminescence signals are lowest at 29°C in antennae, but at peak levels in halteres at the same temperature (Figures S3, S4). Because the *tim-luc* reporter does not encode any TIM protein sequences, we are confident that the bioluminescence levels report endogenous gene expression, and not temperature effects on luciferase activity. In summary, the antennal results confirm the over-compensation phenotypes observed in halteres, thereby strongly suggesting that this is a general inadequacy of peripheral circadian clocks. Moreover, the more severe τ-lengthening defects in antennae point to a correlation of temperature compensation with oscillator strength (see Discussion).

### PER oscillations in brain clock neurons are accurately temperature compensated

To investigate if the over-compensation of peripheral clocks, is also a feature of the neuronal clocks in the fly brain we turned to the *8.0-luc* transgene, where PER expression is restricted to subsets of the dorsal clock neurons (DN and LNd) and excluded from the periphery (Veleri et al., 2003; Yoshii et al., 2009) (Figure S1). Since fruit fly behavioral rhythms are well temperature compensated (Konopka et al., 1989; Singh et al., 2019) and controlled by brain clock neurons (Ewer et al., 1992), we expected *8.0-luc* oscillations to be temperature compensated as well. Indeed, when we recorded *8.0-luc* oscillations in DD, periods were fairly constant between 18°C, 21°C, 25°C, and 29°C with no sign of τ lengthening with increasing temperatures (Q_10_ = 1.06) (Figure 4A, Table 1). Because both *XLG-luc* and *8.0-luc* encode full-length PER-LUC fusion proteins, it seems clear that the observed period-lengthening in peripheral clocks is indeed caused by an impairment of temperature compensation and not by intrinsic properties of PER-LUC fusion proteins.

**Figure 4:**
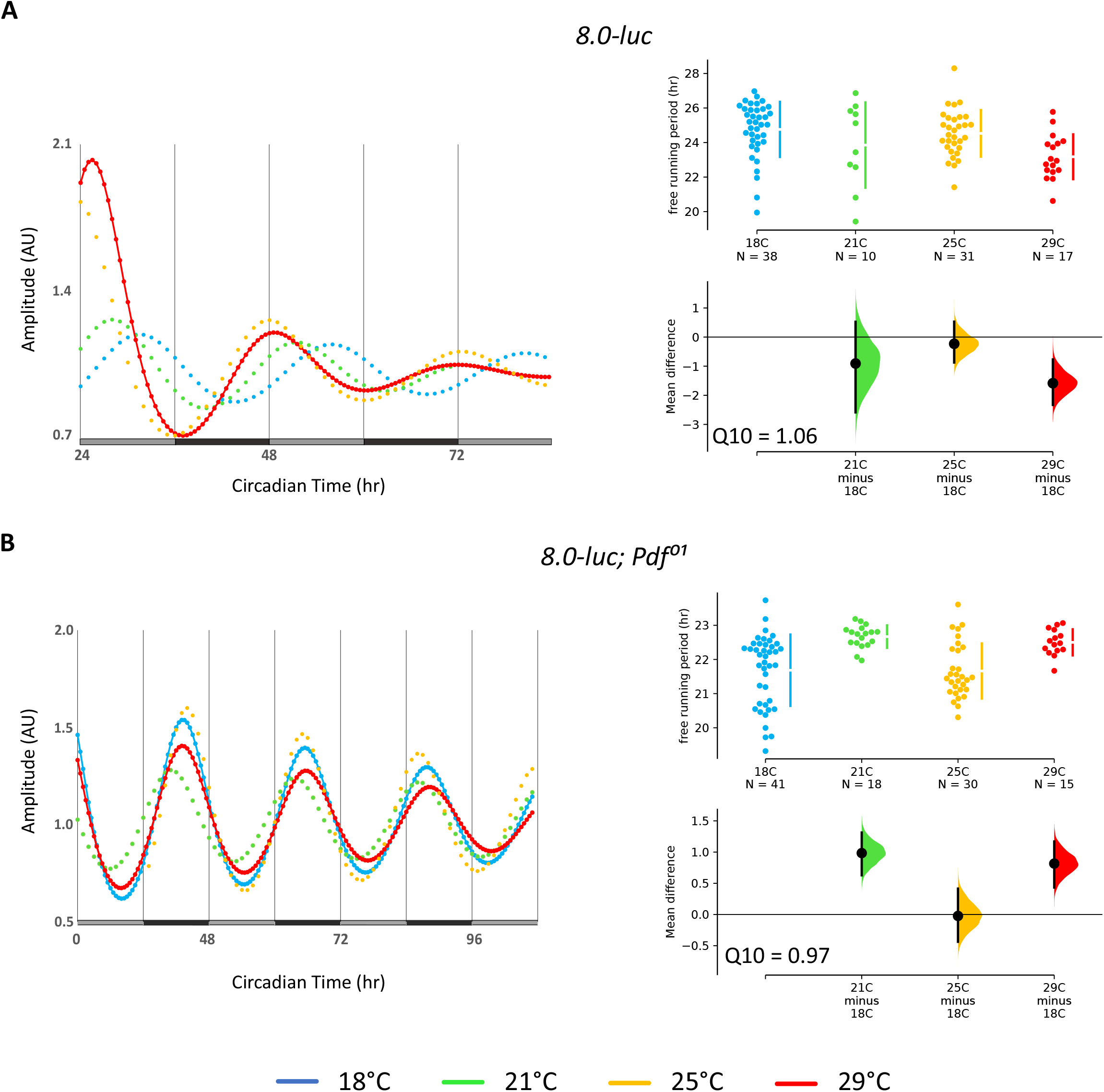
Clock gene expression in central brain clock neurons is temperature compensated and does not require PDF. Bioluminescence recordings of *8.0-luc* flies recorded at different constant temperatures in DD. **(A)** Curve fits of DD data of *8.0-luc* flies (left, and ES comparing individual τ values at different temperatures (right). **(B)** Same as in (A) for *8.0-luc; Pdf*^*01*^ flies.

In contrast to peripheral clocks, the central brain clock in fruit flies is organized in a neuronal network, and these network properties may contribute to accurate temperature compensation. The neuropeptide PDF acts as a synchronizer of this network and differentially influences the period and amplitude of clock protein expression, depending on the particular clock neuronal group (Yoshii et al., 2009). For example, at 25°C and in the absence of PDF, *8.0-luc* oscillations have a short period (22.7 hr) and increased amplitude, most likely reflecting the enhanced PER oscillations in the CRY-negative LNd (Yoshii et al., 2009). To test if PDF is required for temperature compensation, we analyzed bioluminescence rhythms of *8.0-luc Pdf*^*01*^ flies at four different temperatures. As expected, at 25°C we observed robust short-period bioluminescence rhythms, and similar results were obtained for the other three temperatures (Q_10_ = 0.97) (Figure 4B, Table 1). Therefore, PDF is not required for temperature compensation, at least with regard to clock protein expression in the *8.0-luc* expressing neurons.

### Isolated brains maintain slow running, but temperature compensated clock protein oscillations

Brain clock protein oscillations were measured in intact *8.0-luc* flies, and not in isolation as was done for the halteres. To determine if isolated brains also maintain temperature compensated oscillations in DD, we measured bioluminescence rhythms from dissected *tim-TIM-luc* brains in the same culture conditions as described above for the halteres. Surprisingly, oscillations were not as robust as observed in the halteres and the period length was drastically increased to about 30 hr at all 4 temperatures tested (Q_10_ = 0.96) (Figure 5A, B, Table 1). While this indicates that neuronal clocks require some kind of peripheral input in order to run with a 24-hr period, clock protein oscillations in isolated brains are nevertheless properly temperature compensated.

**Figure 5:**
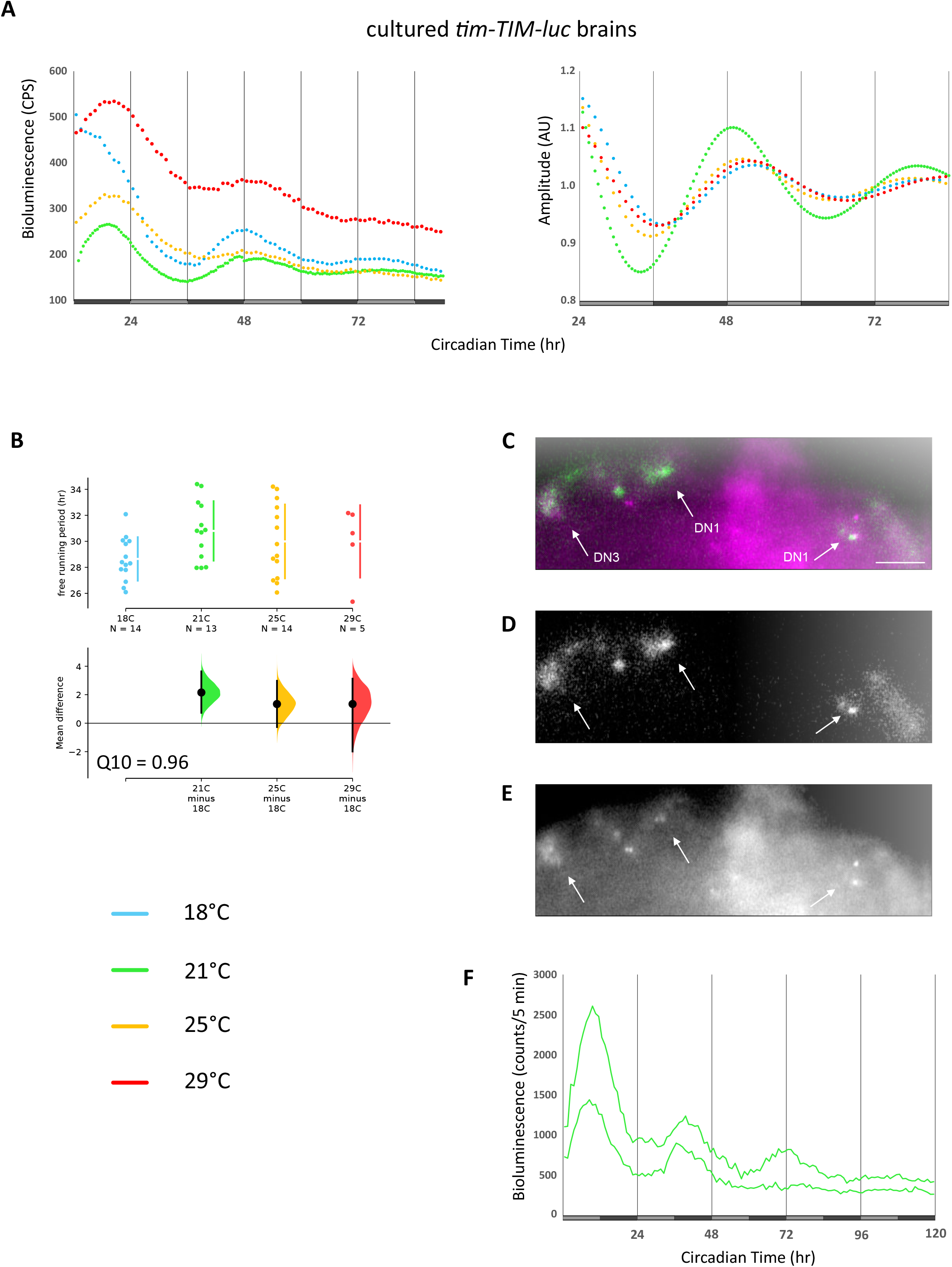
Long-period clock gene expression in cultured brain neurons is temperature compensated. Brains of *tim-TIM-luc* flies were dissected and transferred to cell culture medium followed by bioluminescence recordings in a TopCount plate reader **(A)**, or using a CCD camera **(C-F)**. **(A)** Average raw bioluminescence of brains at the indicated temperature during DD (left), and curve fitted data (right) as described in the legend to Figure 1. N’s are indicated in panel B and Table1. **(B)** ES of individual τ values as described in the legend to Figure 4. **(C-E)** Merged (C), bioluminescence (D, green in merge), and fluorescence images (E, magenta in merge) of *tim-TIM-TOMATO*; *tim-TIM-luc* brains. Scale bar = 50µm. **(F)** CCD camera bioluminescence recordings of the two groups of DN1 neurons shown in panels C-E at 21°C in DD (see Transparent Methods). For τ see Table 1.

The bioluminescence signals recorded from isolated *tim-TIM-luc* brains are expected to be composed of clock glia cells in addition to the clock neurons (Ewer et al., 1992). Since the TopCount plate-reader measures overall light-output from individual wells, it does not allow for any spatial resolution of gene expression. To confirm if clock neurons in culture indeed oscillate with long ~30 h periods, we therefore applied a bioluminescence imaging system (LV200, Olympus) to measure clock gene oscillations in a subset of the clock neurons. For this *tim-TIM-luc* flies were combined with a red-fluorescent reporter, to label the nuclei of all clock neurons (Mezan et al., 2016). Brains of *tim-TIM-TOMATO*; *tim-TIM-luc* flies were dissected and using the fluorescence mode of the microscope, we focused on a subset of the DN1 clock neurons (Figure 5C, E). In the absence of fluorescence light (Figure 5D), photons were then captured automatically once per hour for 4 days using a cooled CCD camera (see Transparent Methods). Within 2 DN1 clusters bioluminescence oscillated with a period around 31 h (Figure 5F, Movie S1, Table 1). Combined, the brain culture results indicate that clock neurons in isolated brains oscillate with a period ≥ 30 h and that these oscillations are temperature compensated.

## Discussion

We report here important characteristics of peripheral clocks in the insect *Drosophila melanogaster*. While previous work showed that these peripheral clocks can be synchronized tissue-autonomously to LD and temperature cycles, it was not clear if they indeed constitute self-sustained and temperature compensated oscillators. Hence, up to now it was questionable if peripheral fly clocks qualify as true circadian clocks. Our work shows, that at least the haltere peripheral clock maintains robust and self-sustained 24-hr oscillations, and that the period of PER and TIM protein expression rhythms increases with temperature (over-compensation). Transcriptional rhythms for both genes are less over-compensated, with Q_10_ values close to 1 (0.90 for *tim-luc* and 0.94 for *plo*). While this may indicate better temperature compensation for transcription, in the antannae we observed more severe over—compensation with the transcriptional reporters compared to the protein reporters (Figure 3, Table S1). In summary our results suggest that over–compensation in the face of increasing ambient temperatures is a general feature of peripheral clocks in fruit flies, while the molecular mechanisms of temperature compensation may differ between different tissues.

### Differences between central and peripheral oscillators

Why are peripheral clocks not able to maintain stable period lengths’ at different temperatures? One possible answer is the different make-up of the molecular oscillator in peripheral clocks, which requires CRY function in most studied cases (Ivanchenko et al., 2001; Krishnan et al., 2001; Levine et al., 2002). One exception is the cuticle deposition rhythm, but it is not known if this rhythm is temperature compensated (Ito et al., 2008). Another possibility is related to the cell-type harboring central versus peripheral clocks. In addition to rhythms of clock gene expression, central clock neurons also display clock regulated daily rhythms of neuronal activity, which is thought to feedback and enhance clock gene expression rhythms (reviewed in (Allen et al., 2017)). Considering the non-neuronal nature of a subset of the peripheral clock cells (e.g., MT, testes, epidermal cells that give rise to the cuticle deposition rhythm) it is possible that neuronal network properties of the circadian clock neurons contribute to temperature compensation. For example, in the mammalian Suprachiasmatic nuclei (SCN), coupling between the individual clock neurons prevents their synchronization to temperature cycles, suggesting that network properties contribute to resilience against temperature induced changes to the molecular clock (Abraham et al., 2010; Buhr et al., 2010). Our finding that removal of the circadian neuropeptide PDF, a potent coupling signal between different circadian clock neurons, did not impair temperature compensation of clock neurons, argues against an important role of network communication in temperature compensation. Nevertheless, our results do not rule out that PDF-independent neuronal network properties contribute to temperature compensation of the fruit fly brain clock.

Alternatively, the temperature-dependence of peripheral clock oscillations may contribute to the temperature-independence of the brain oscillators. Perhaps the brain clocks are connected to the peripheral clocks and somehow receive temperature information encoded by the period-changes in the peripheral oscillators. This way, the central oscillators could be instructed to speed up or slow down their molecular oscillations, in order to maintain a stable τ at different temperatures. In this scenario, the ‘outside’ location of the peripheral oscillators (antennae, haltere, legs) would help them to react quickly to temperature changes. Although this is an interesting speculation, the fact that we observed temperature compensated rhythms of ≥ 30 hr in isolated brains, shows that peripheral clocks are not required for temperature compensation of the brain oscillator. We have no explanation for why clock gene oscillations in isolated brains show long periods, but the reproducible results, obtained with different recording methods, indicate that they are genuinely long. In fact, long periods of *tim-luc* expression in subsets of the clock neurons of isolated brains, recorded with a similar CCD camera system were reported previously (Sellix et al., 2010). One reason for the long periods could be that brains in culture change their neuronal firing properties. For example, it has been shown that the bursting firing mode of the l-LNv decays rapidly within the first 30 min after dissection and also depends on input from the visual system (Muraro and Ceriani, 2015). It is therefore likely that the firing properties of the clock neurons after several days in culture are quite different compared to an intact brain. Because neuronal firing feeds back on molecular oscillations, changed neuronal activity may influence (slow down) the free running molecular oscillations independent of temperature. Future work will reveal if the long periods are linked to specific subsets of clock neurons or a general feature observed in isolated brains in culture. The latter would indicate that the period of central clock neurons is normally accelerated by an unknown input from outside the brain, as for example the visual system or other peripheral clock containing tissues and sensory organs.

### Robustness of peripheral and central oscillators

Our comparison between peripheral oscillators revealed that the haltere oscillator is more robust and also less over-compensated compared to the antennal oscillator. It is possible that this correlation reflects a causal relationship between oscillator strength and temperature compensation. On the other hand, although the haltere oscillator is very robust and basically non-dampening (Figure 1A), it does show significant over-compensation compared to the brain oscillator, ruling out that oscillator strength is the unique determinant for fully functional temperature compensation.

### Why a peripheral clock in the haltere?

Although it is formally possible that the haltere clock supports temperature synchronization of the central brain clock (see above), it seems more likely that it fulfils a more direct physiological function in this proprioceptive sensory organ. Clock gene expression in the *Drosophila* haltere was observed in cells that most likely represent campaniform sensilla, the principle haltere mechanosensory neurons (Sehadova et al., 2009; Yarger and Fox, 2016). In contrast, no clock gene expression in the other haltere mechanosensory cell type, the chordotonal organ neuron, could be detected (Sehadova et al., 2009). We therefore conclude that the rhythmic haltere gene expression we report here, emanates from the campaniform sensilla. At present, it is unknown which biological rhythm(s) the haltere clock regulates. Compared to other peripheral clocks the haltere rhythms are strikingly more robust (e.g., compare Figures 1, 2, S3 to Figures 3, S4), suggesting that the haltere clock does have an important physiological function.

## Conclusions

In summary, our work reveals surprising features of peripheral clocks, based on the discovery of a robust and non-dampening 24-hr oscillator in the fly haltere. Unlike the central clock neurons in the brain, the haltere oscillator is over-compensated against temperature increases. Our findings extend beyond poikilothermic invertebrates, because it has been reported that peripheral clocks in both ectothermic and endothermic vertebrates are also over-compensated (Barrett and Takahashi, 1995; Izumo et al., 2003; Kaneko et al., 2006; Lahiri et al., 2005; Reyes et al., 2008; Tsuchiya et al., 2003). Careful comparison of our invertebrate data with the vertebrate studies reveals that in embryonic zebra fish,- and mammalian fibroblast cell lines, as well as in dispersed chick pineal cells, clock gene oscillations slow down with increasing temperature, with similar Q_10_ values as reported here (Q_10_ ≤0.9) (Izumo et al., 2003; Lahiri et al., 2005; Tsuchiya et al., 2003). Circadian clocks of various cultured adult zebra fish tissues have Q_10_ values between 0.9 and 1.0 (Kaneko et al., 2006), while in cultured tissues of *mPer2-luc* mice the only tissue with a Q_10_ ≥ 1.0 was the SCN, and all peripheral clocks slowed down with temperature. Moreover, the liver oscillator did not show self-sustained oscillations below 37°C (Reyes et al., 2008) similar to what we show here for the antennae at 29°C. Combined, the available data from various circadian model organisms point to a more prominent difference between the make-up of peripheral and central circadian oscillators than previously thought. This is also supported by the lack of a PRC dead-zone in mammalian peripheral clocks (Balsalobre et al., 2000). Our results indicate that in *Drosophila* the differences between central (Q_10_ = 1) and peripheral clocks are particularly prominent, as Q_10_ values for PER and TIM protein rhythms in the haltere and transcriptional and protein rhythms in the antennae are at the lower end, or even lower, compared to what has been reported for most vertebrate peripheral tissues (Table 1 and S1). The question which of the differences between peripheral and central oscillators (e.g., molecular and/or neuronal network properties) is responsible for the different temperature compensation properties cannot be finally answered, and requires more detailed analysis and comparisons between different organisms, tissues, and organs.

## Acknowledgements

We thank Patrick Emery and Sebastian Kadener for fly stocks. We thank Luis Garcia for help with statistics, graphical design, and discussions, and Mechthild Rosing and Carolina Camelo for technical support. We particularly thank Baris Can Mandaci for his preliminary work leading up to this study. This work was funded by a grant from the Deutsche Forschungsgemeinschaft given to RS (STA421/7-1).

---

**Movie S1: Bioluminescence recordings of two DN1 clusters during constant conditions.** DN1 clusters were identified with *tim-TIM-TOMATO* fluorescence and bioluminescence was recorded using an EM-CCD camera for 5 consecutive days in DD and 21°C. See text and Figure 5C-F for details.

