## Supplemental Data for "A robust and self-sustained peripheral circadian oscillator reveals differences in temperature compensation properties with central brain clocks"

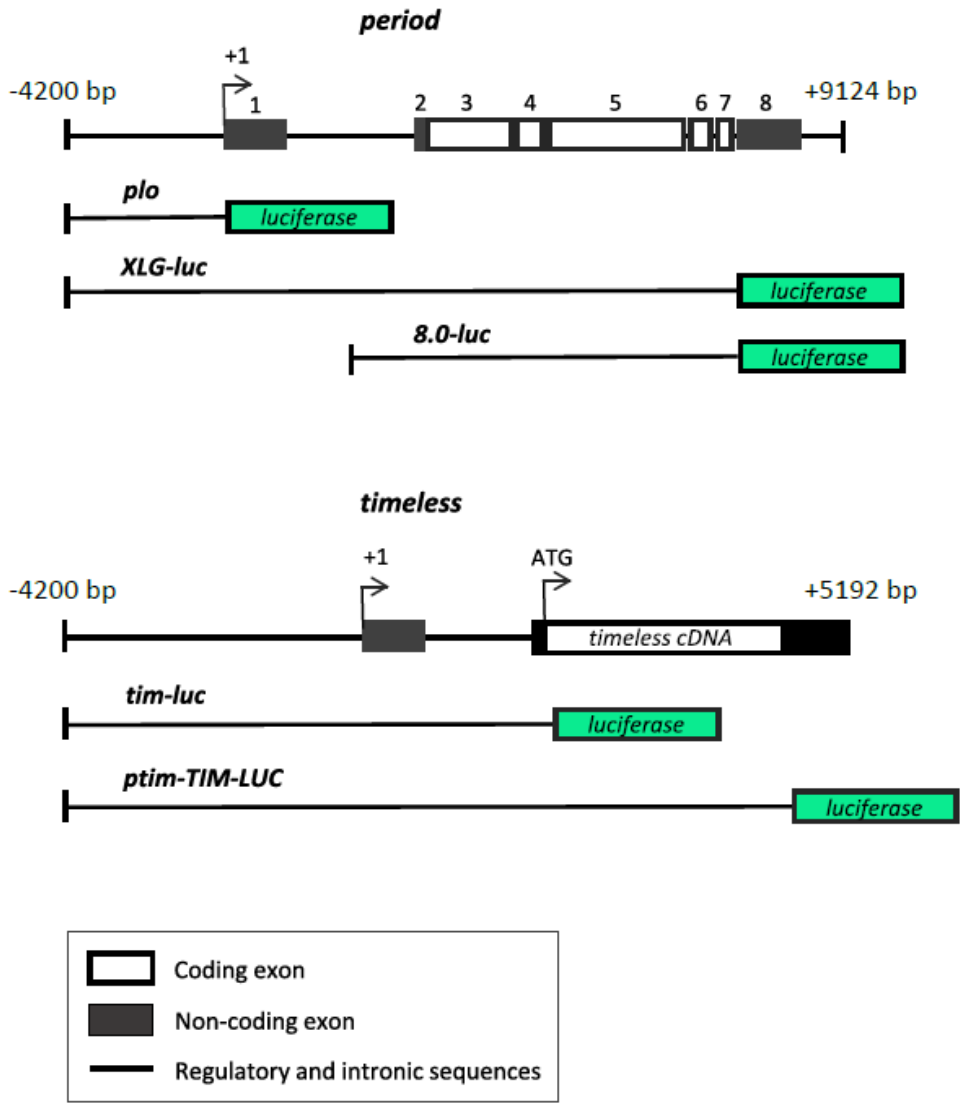

**Figure S1: Map of the *period* and *timeless* luciferase constructs, Related to Figures 1-5.** Top: genomic organization of the *per* locus and *per* content of the 3 *per-luciferase* constructs applied here. *plo* contains only 5’ *per-*regulatory sequences, reporting *per* transcription (Brandes et al., 1996; Stanewsky et al., 1997), while *XLG-luc* and *8.0-luc* report PER protein expression (Veleri et al., 2003). Due to the lack of regulatory *per* promoter sequences, *8.0-luc* expression is restricted to subsets of the clock neurons and excluded from the peripheral clocks (Gentile et al., 2013; Veleri et al., 2003; Yoshii et al., 2009). Bottom: map of a *timeless* construct containing ~6.3 kb 5’-regulatory sequences fused to the *tim* cDNA promoter (Rutila et al., 1998) and the extend of *tim* sequences contained in the *tim-luc* and *tim-TIM-luc*, reporting *tim* transcription and TIM protein expression, respectively (Lamba et al., 2018; Stanewsky et al., 2002).

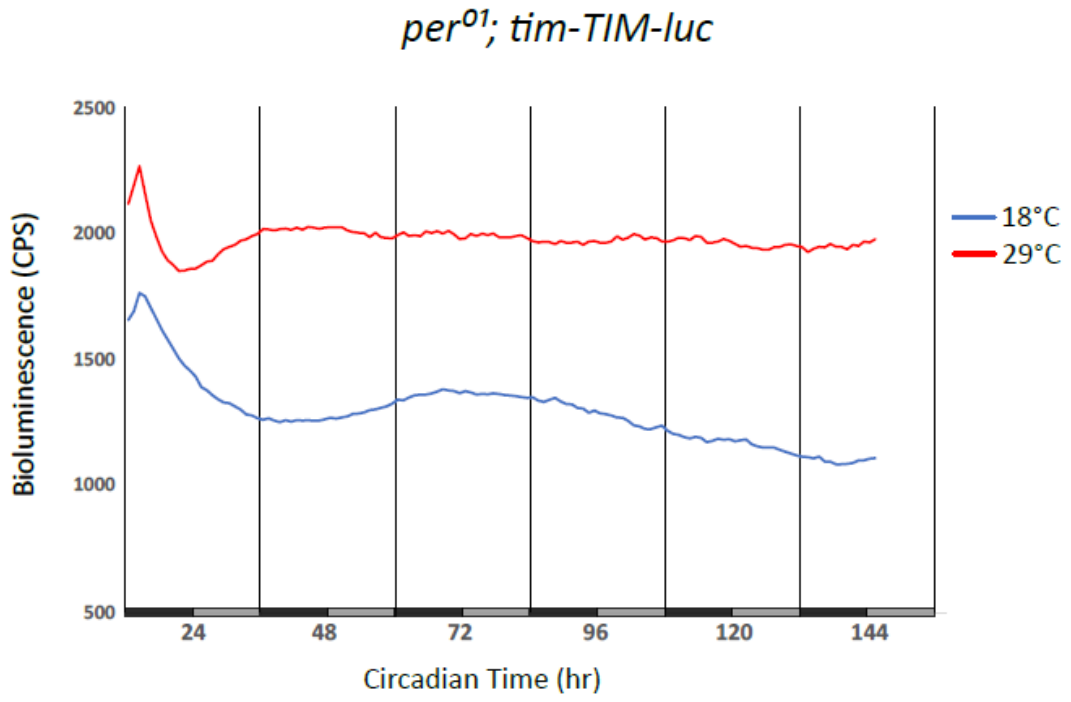

**Figure S2: Rhythmic *tim-TIM-luc* expression in the haltere depends on a functional clock, Related to Figure 1.** Bioluminescence recordings of halteres dissected from *per^01^; tim-TIM-luc* flies during DD at 18°C (n=48) and 29°C (n=46). Note the arrhythmic, but drastically increased (> 10 fold) expression levels compare to *per^+^* due the lack of PER repressor activity (compare to Figure 1).

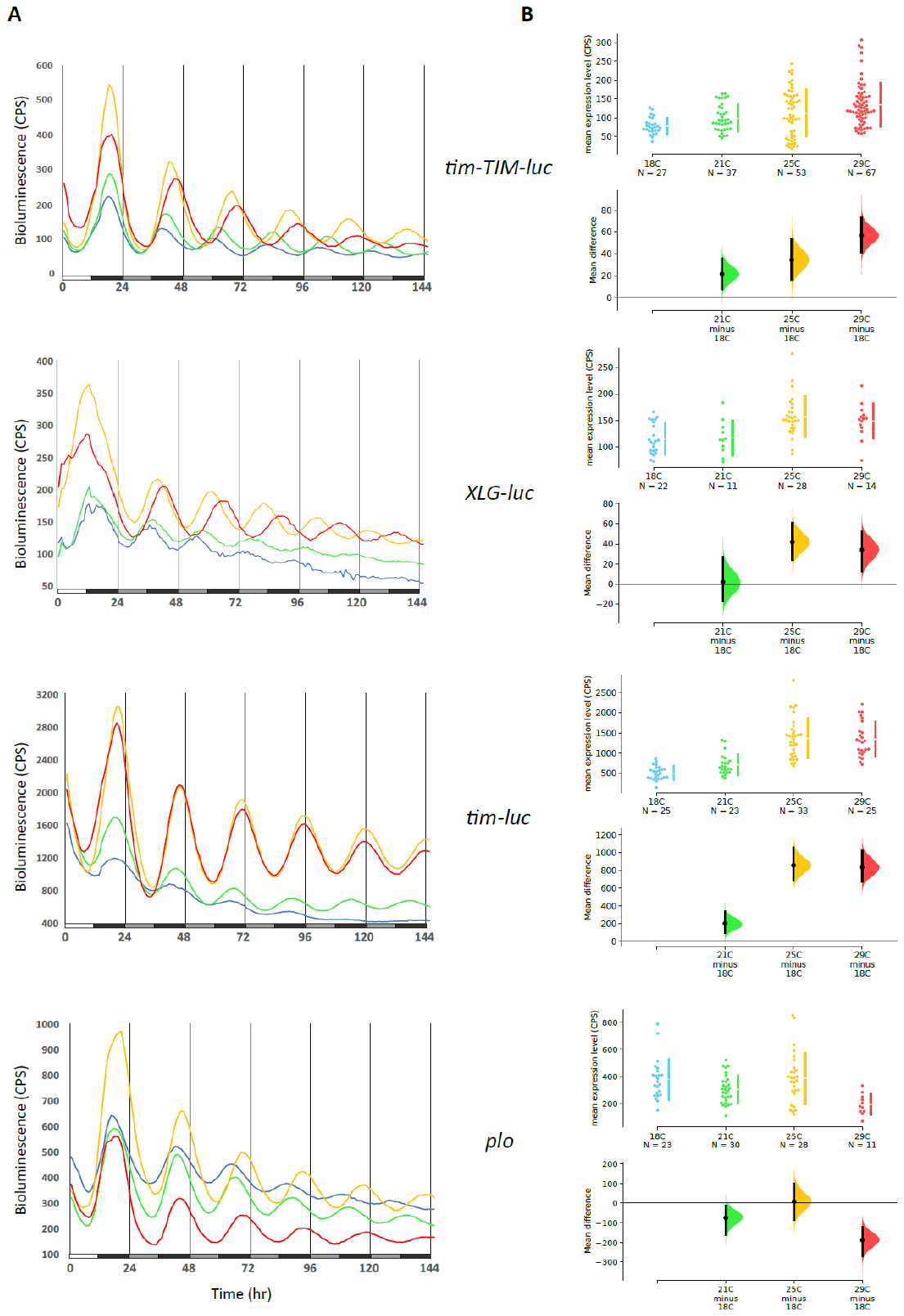

**Figure S3: Raw levels of reporter gene expression in halteres at different temperatures, Related to Figures 1, 2, Table 1. (A)** Bioluminescence recordings of halteres dissected from *tim-TIM-luc* (replotted from Figure 1B), *XLG-luc*, *tim-luc*, and *plo* flies at the indicated constant temperatures. The plots show raw-data averages from the last day in LD followed by 5 days in DD. White bars on the X-axis indicate lights-on, black bars lights-off, and grey bars, subjective day. **(B)** Gene expression levels reflecting the mean CPS of a single haltere pair during the entire recording. To precisely visualize potential gene expression differences between temperatures we applied ES (see legend to Figure 1D and Transparent Methods for details).

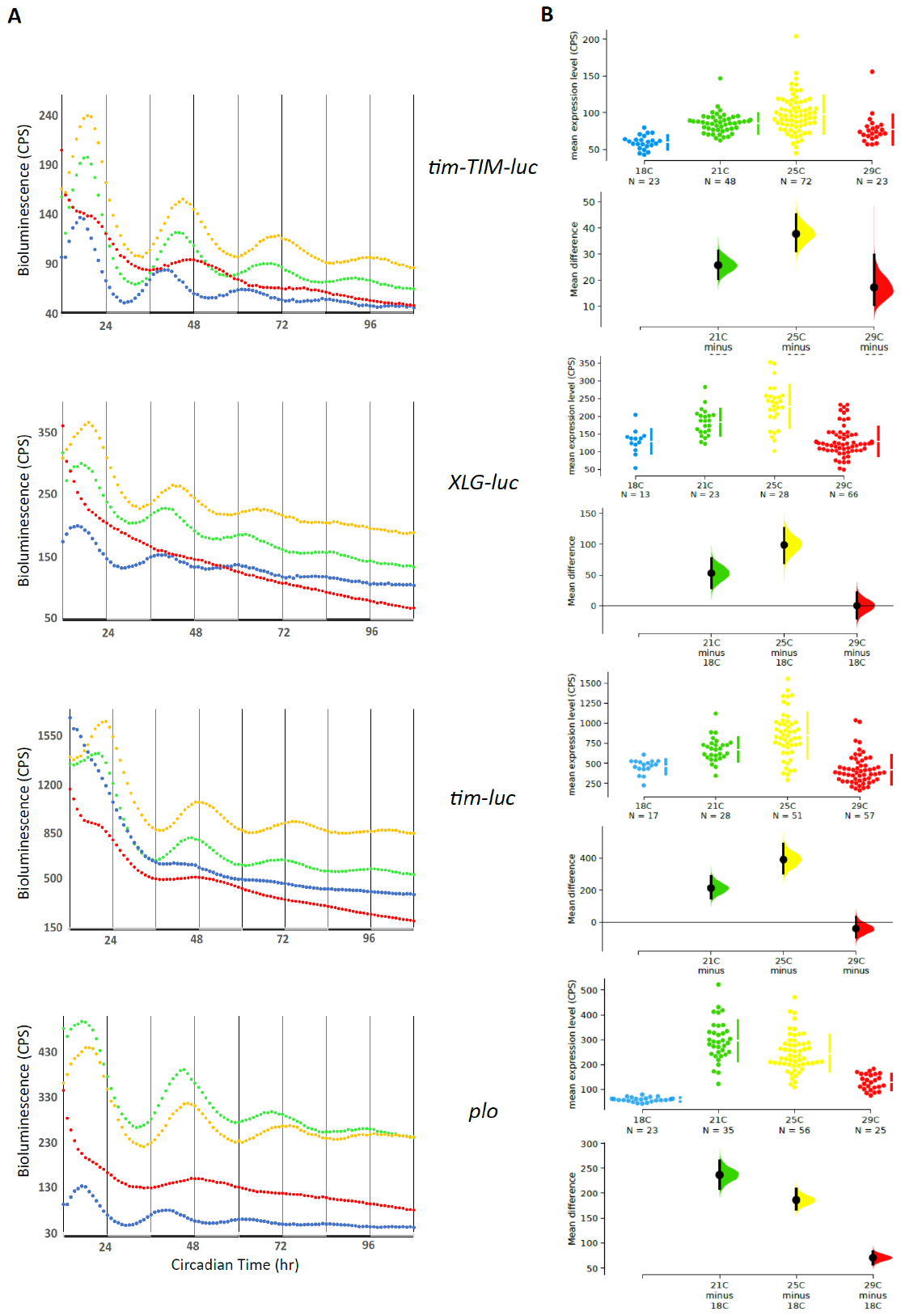

**Figure S4: Raw levels of reporter gene expression in antennae at different temperatures, Related to Figure 3 and Table S1. (A)** Bioluminescence recordings of antennae dissected from *tim-TIM-luc,* *XLG-luc*, *tim-luc*, and *plo* flies at the indicated constant temperatures. The plots on the left show raw-data averages from the 4 days in DD immediately following LD entrainment. Black bars indicate subjective night; grey bars subjective day. **(B)** Gene expression levels reflecting the mean CPS of a single antennal pair during the entire recording. To precisely visualize potential gene expression differences between temperatures we applied ES (see legend to Figure 1D and Transparent Methods for details).

**Table S1: Free running period (τ) and expression levels of clock gene expression in antennae, Related to Figure 3 and S4.**

| Genotype | °C | N | Tau ± S.E.M (h) | Error ± S.E.M | Mean ± S.E.M (CPS) |
| --- | --- | --- | --- | --- | --- |
| **antennae** |  |  |  |  |  |
| *plo* | 18 | 23 | 22.6 ± 0.1 | 0.10 ± 0.01 | 59.7 ± 1.9 |
|  | 21 | 35 | 26.4 ± 0.2 | 0.09 ± 0.01 | 296.3 ± 13.8 |
|  | 25 | 56 | 27.2 ± 0.1 | 0.10 ± 0.01 | 245.8 ± 9.9 |
|  | 29 | 90 (25) | 33.9 ± 0.5 | 0.39 ± 0.02 | 102.3 ± 3.6 |
|  |  |  | ***Q_10_ = 0.69*** |  |  |
| *tim-luc* | 18 | 47 (17) | 21.0 ± 0.5 | 0.43 ± 0.02 | 482.1 ± 17.0 |
|  | 21 | 28 | 26.1 ± 0.1 | 0.07 ± 0.01 | 673.3 ± 28.7 |
|  | 25 | 51 | 27.3 ± 0.1 | 0.06 ± 0.01 | 851.4 ± 40.3 |
|  | 29 | 83 (58) | 33.7 ± 0.1 | 0.29 ± 0.02 | 395.7 ± 20.2 |
|  |  |  | ***Q_10_ = 0.65*** |  |  |
| *XLG-luc* | 18 | 13 | 22.8 ± 0.3 | 0.25 ± 0.02 | 129.2 ± 9.7 |
|  | 21 | 23 | 22.7 ± 0.2 | 0.19 ± 0.02 | 182.3 ± 7.9 |
|  | 25 | 36 | 24.1 ± 0.2 | 0.16 ± 0.01 | 232.2 ± 9.3 |
|  | 29 | 66 | AR | N/A | 129.5 ± 5.1 |
|  |  |  | ***Q_10_ = 0.92*** |  |  |
| *ptim-TIM-luc* | 18 | 23 | 22.6 ± 0.1 | 0.10 ± 0.01 | 59.7 ± 1.9 |
|  | 21 | 48 | 25.4 ± 0.1 | 0.06 ± 0.01 | 85.6 ± 2.0 |
|  | 25 | 72 | 26.7 ± 0.1 | 0.09 ± 0.01 | 97.6 ± 3.0 |
|  | 29 | 57 (23) | 31.8 ± 0.1 | 0.35 ± 0.02 | 67.0 ± 2.3 |
|  |  |  | ***Q_10_ = 0.73*** |  |  |

See Table 1 legend for details. Numbers in parentheses indicate N used for calculation of τ in cases where not all samples were rhythmic (cut-off error < 0.5). Q_10_ values were calculated using the τ values for 18°C and 29°C except for *XLG-luc* (18°C and 25°C).

**Transparent Methods**

**Flies**

Flies were kept at 25°C or 18°C on common cornmeal-yeast-sucrose food under light:dark cycles and ~60% humidity. *plo*, *tim-luc, XLG-luc, tim-TIM-luc* are *per* or *tim* fusions with the firefly *luciferase* cDNA that were all described previously and shown to report transcriptional or protein rhythms of the two clock genes in peripheral tissues (Brandes et al., 1996; Lamba et al., 2018; Stanewsky et al., 2002, 1997; Veleri et al., 2003). The transgenic *8.0-luc* line was described previously and encodes the entire PER protein fused to the firefly Luciferase cDNA (Veleri et al., 2003). Due to the absence of the *per* promoter, the 1^st^ non-coding exon, and large parts of the 1^st^ intron, reporter expression is limited to subsets of the DN1-3 and LNd both during TC and LD cycles (Gentile et al., 2013; Veleri et al., 2003; Yoshii et al., 2009). *Pdf^01^* (Renn et al., 1999) and *per^01^* (Konopka and Benzer, 1971) loss-of-function mutants were combined with the respective reporter constructs using standard crosses. The *tim-TIM-TOMATO* (chromosome *2)* flies express a destabilized red fluorescence protein in all clock cells (Mezan et al., 2016) and were combined with *tim-TIM-luc* (chromosome *3)* using standard crosses.

**Bioluminescence measurements of adult flies**

Luciferase expression of individual flies carrying the *8.0-luc* *period-luciferase* reporter gene was measured as described in (Veleri et al., 2003). 3-4 day old males were loaded in 96-well microtiter plates containing 100ul of 5% sucrose, 1% agar and 15mM luciferin. Bioluminescence was detected with a TopCount Multiplate Reader (Perkin Elmer) for serval days during conditions of the designated constant temperature and DD after the flies were entrained to LD at the same temperature for 2 days. Bioluminescence was measured once or twice per hour and data were analyzed using ChronoStar software (Klemz et al., 2017).

**Culturing and plate reader bioluminescence measurements of *Drosophila* peripheral tissues and brains**

Halteres and antennae of two- to seven-day old transgenic *plo*, *XLG-luc*, *tim-luc* or *ptim-TIM-luc* males kept under 12 h :12 h LD at 25°C were bilaterally dry dissected. Each pair was transferred into every other well of a 96 well plate (Topcount, Perkin Elmer) filled with culture medium containing 80% Schneider’s medium (Sigma-Aldrich), 20% inactivated Fetal Bovine Serum (Capricorn Scientific, Ebsdorfergrund, Germany) and 1% PenStrep (Sigma-Aldrich) and fortified with Luciferin (Biosynth AG, Switzerland) to a final concentration of 226 µM. Plates were sealed with clear adhesive covers and transferred to a TopCount plate reader (Perkin Elmer). Bioluminescence emanating from each well was measured hourly or half-hourly in LD for two days, followed by 5 days of constant darkness (DD) at 18°C, 21°C, 25°C or 29°C, respectively, as described above for intact *8.0-luc* flies. For brain cultures, dissections of *ptim-TIM-LUC* male brains were done in Ca^2+-^free Ringer’s solution after briefly being rinsed in a 96% EtOH solution. Single brains were placed into a well containing the same medium as described above, but supplemented with 1 mM Luciferin.

**Bioluminescence imaging of adult brains using the LV200 system**

*ptim-TIM-TOMATO; ptim-TIM-luc* flies were synchronized to LD cycles at 21°C for at least 3 days. Thereafter, brains were dissected and collected in Ca^2+^ free Ringer’s solution within 1 hour and subsequently placed on a Cellview glass bottom imaging dish (35x10mm; Greiner Bio One) treated with heptane glue. Brains were placed into a droplet of Ca^2+^ free Ringer’s solution that was placed on top of the glue. The solution was removed leaving the brains glued to the dish and immediately replaced by a drop of culture medium (see above) containing 1 mM Luciferin where after additional culture medium was carefully added. Culture dishes were then placed into a sterile transparent glass container and entrained for two days in LD at 21°C, replacing two thirds of culture medium with fresh one each day. At day 3, culture dishes were transferred to a LV200 (Olympus) microscope placed in a temperature controlled dark room. Positioning of the brains was done by means of a mobile stage equipped with a Stage Top Incubator (Tokai Hit, Japan) and bright-field illumination. Fluorescence and use of the *ptim-TIM-TOMATO* construct allowed focussing on individual DN1 clock neuron cell bodies based on anatomical position using 40X magnification (Figure 4C-E). Bioluminescence was then measured during 5 min exposure periods with an EM gain of 400 (0.688 MHz EM-CCD camera CAM-ImageEM X2, Hamamatsu, Japan) followed by 5 s exposure to fluorescence light (to control focus) with an EM gain of 4. Exposures were performed once every hour from ZT12 onwards in DD for 5 days generating 512x512 pixel movies (see Movie S1). Images were processed with the CellSense software (Olympus, Japan). Average luminescence intensity of closely grouped DN1 cell bodies was measured within manually defined regions of interest (ROIs) over time, based on *ptim-TIM-TOMATO* signals. Dynamic ROIs were determined by re-positioning ROIs frame to frame to account for flattening of the cultured brains and to exclude cosmic rays as described previously (Roberts et al., 2015). Background correction was performed by subtracting the intensities of equally sized ROI_background_ from ROI_cell bodies_ . Raw bioluminescence data were further processed with ChronoStar software to calculate period length as described above. Single frame images for Figure 6 C-E were processed in Fiji using a 1:14 gamma correction for the fluorescence channel.

**Period and expression level calculation of bioluminescence data using ChronoStar software**

Using ChronoStar software (Klemz et al., 2017) raw data from individual wells (tissues or flies) were first detrendend, using a running average with a 24 h window. After subtracting the resulting trend line from the raw data, the DD part of the data are subjected to a sinus fit operation. The parameters of the resulting curve include a period estimate (Tau in hr), along with a dimension less ‘Error’ value, depicting the correlation between the curve fit and the detrended data (1 minus correlation coefficient, so the lower the error value the more trustable the period value). For this study, we applied a cut-off of <0.5 for the error value for all tissue culture experiments and <0.7 for the experiments with adult flies and only samples meeting these criteria were included in the average calculations for ‘Tau’ and ‘Error’ plotted in Table 1. In addition, ChronoStar determined the mean expression level for the entire time series of each well, and the average ‘Mean’ is also tabulated in Table 1. Furthermore, the individual values for ‘Tau’ and ‘Mean’ were used for the estimation graphics and statistical tests (see below). To generate line and scatter graphs raw and curve fitted averaged time series were plotted in Excel and figures generated in Inkscape.

**Estimation graphics and statistical tests**

For the statistical analysis of the data, estimation statistics has been used. This approach gives a more informative way to analyze and interpret results (Claridge-Chang and Assam, 2016). It focuses on the effect size of one's experiment, as opposed to significance testing. While significance testing (p-values) focus on the acceptance or rejection of the null hypothesis, estimation stats focus on the magnitude of the effect size (i.e. mean difference) and its precision (Ho et al., 2019).

Data were analysed using DABEST (Ho et al., 2019), using the website available under https://www.estimationstats.com/#/. In order to compare the different groups of data at the same time, providing an overall view, a shared-control plot was used, which is analogous to an ANOVA with multiple comparisons (Ho et al., 2019). This plot is divided in two parts. First, the top part shows all the observed data points, showing the underlying distribution, right to every data set the mean and standard error are plotted as a discontinuous line, the gap indicates the observed mean. The bottom part shows the effect size (mean difference) as a bootstrap 95% confidence interval (CI) with BCa correction (Efron, 1979). Here the magnitude of the mean differences between groups can be easily compared with the reference. For all our plots the most left data set (18°C) is considered the reference and all mean differences are plotted relative to it. The size of the CI shows the precision of the mean difference. The smaller it is the more confident is the measure. The further apart the mean difference (black dot inside CI) is from the reference line, the more difference exists between the datasets (p value < 0.05). If the CI is cutting the reference line then, both datasets may originate from the same distribution. Therefore, they are not truly different (p value > 0.05).
